# Interactions between saccades and smooth pursuit eye movements in marmosets

**DOI:** 10.1101/2024.01.07.574533

**Authors:** Jagruti J. Pattadkal, Carrie Barr, Nicholas J. Priebe

## Abstract

Animals use a combination of eye movements to track moving objects. These different eye movements need to be coordinated for successful tracking, requiring interactions between the systems involved. Here, we study the interaction between the saccadic and smooth pursuit eye movement systems in marmosets. Using a single target pursuit task, we show that saccades cause an enhancement in pursuit following a saccade. Using a two-target pursuit task, we show that this enhancement in pursuit is selective towards the motion of the target selected by the saccade, irrespective of any biases in pursuit prior to the saccade. These experiments highlight the similarities in the functioning of saccadic and smooth pursuit eye movement systems across primates.

**SIGNIFICANCE STATEMENT:** We study the coordination between the smooth-pursuit and saccadic eye movement systems in marmosets using single and multiple object motions. We find that saccade to a target increases pursuit velocity towards the target. If multiple objects are visible, saccade choice makes pursuit more selective towards the saccade target. Our results show that coordination between different eye movement systems to successfully track moving objects is similar between marmosets and primates.

## INTRODUCTION

Primates use a combination of eye movements to visually track objects in our environment (Rashbass, 1961). Saccadic eye movements bring objects of interest onto the fovea whereas smooth pursuit eye movements stabilize moving objects onto the retina. These two eye movements are sensitive to distinct aspects of object information: saccades are primarily sensitive to object position, and pursuit eye movements are primarily sensitive to object motion (Rashbass, 1961). While these two classes of eye movements are distinct, the retinal input and scene interpretation driving these movements are shared (Krauzlis et al., 1997; Gardner and Lisberger, 2001; Adler et al., 2002; Gardner and Lisberger, 2002; Krauzlis and Dill, 2002; Liston and Krauzlis, 2003; Carello and Krauzlis, 2004; Krauzlis, 2004; Liston and Krauzlis, 2005; Schoppik and Lisberger, 2006; Case and Ferrera, 2007). Achieving the common goal of selecting and tracking visual objects thus requires the coordination of both saccades and smooth pursuit eye movements.

Here we examined how saccades alter pursuit eye movements in situations in which marmosets track either single or multiple targets. While the primary drive to pursuit eye movements is target motion, it has long been recognized that pursuit is susceptible to extra-retinal factors (Schwartz and Lisberger, 1994; Ferrera and Lisberger, 1995). These factors, including a selection of a target, influence the direction of pursuit eye movements (Stone et al., 1996; Beutter and Stone, 1998) and the degree to which the pursuit system is engaged. Saccadic eye movements to a target, potentially reflecting target selection, lead to an increase in the amplitude of smooth pursuit in humans (Robinson, 1965; Luebke and Robinson, 1988) and macaques (Lisberger, 1998). This post-saccadic enhancement has been interpreted as a reflection of the engagement of the pursuit system (Schwartz and Lisberger, 1994; Churchland and Lisberger, 2002) when a target has been selected. Evidence for this engagement also exists in marmosets: responses to a target perturbation during pursuit are larger than the same target perturbation from fixation (Mitchell et al., 2015). It is unclear, however, if this engagement is related to the saccadic system, as these perturbations were made during steady state pursuit.

The visual world is often composed of multiple moving objects. To selectively track a single moving object, animals generally saccade to an object, to reduce the positional error and then pursue that object to reduce the velocity error. As such, there is a need for coordination between the two eye movements to ensure that the saccade and pursuit match the position and velocity of the selected target. Previous studies in humans (Krauzlis et al., 1999) and macaques (Gardner and Lisberger, 2001) have shown that target selection decisions for saccade and pursuit are indeed linked, but it remains unclear whether these systems are working in parallel or serial. Here we demonstrate that, as in macaques and humans, saccades influence the gain of pursuit eye movements. We also find, however, that smooth eye movements exhibit information about target selection prior to saccadic eye movements, indicating that choice about pursuing an object does not require saccadic eye movements.

## MATERIALS AND METHODS

Two male common marmosets were used for this study. The animals were surgically implanted with a titanium headpost, attached to the skull using Metabond (Parkell). Post recovery, they were trained to be head-fixed and perform fixation and pursuit (Mitchell et al., 2015; Pattadkal et al., 2023a). To motivate animals for behavioral tasks, they were on food control with weights maintained at 85-90% of baseline weight. Baseline weight was calculated as average weight during a 4-week period each year when the animal is on free food and water and no behavioral training occurs. All procedures conformed to National Institutes of Health guidelines and the guidelines of the University of Texas at Austin Institutional Animal Care and Use Committee.

We used the Eyelink 1000 plus system for IR eye tracking. The data was collected in pupil-CR mode at 1kHz rate. Stimuli were displayed on FlexScan T761 50 cm (19 inch) Class Color Display cathode ray tube monitor with a refresh rate of 85 Hz. The distance between monitor and the animal is 50 cm with a field of view was 35° by 28°. Stimuli were shown on a gray background with mean luminance 31.5 cd/m2. The maestro software was used to control the behavioral tasks and collect data. (https://sites.google.com/a/srscicomp.com/maestro/).

Tasks started with animals fixating at a target presented at the center of the screen for 500-1000 ms and 200-500 ms grace period. Marmoset faces or small dots were used as targets. For single target tasks, the target then stepped either in the direction (positive step) or in the opposite direction (negative step) of upcoming target motion. The target then moved at a constant speed for 700-800 ms. For a successful trial, animals had to follow the moving target, remaining within a fixation window of 2 deg. For animal 1, target moved at 5 deg/s and step size was +1 or -1 deg. Target directions with either rightwards (0), upwards (90) or leftwards (180). For animal 2, target moved at 8 deg/s and step size was of -1.6 deg. Target directions with either rightwards (0), upwards (90), leftwards (180) or downwards (270). For two target tasks, after the initial fixation, two targets appeared and made a step jump. Both targets then moved in their respective direction for up to 800 ms. After a variable period, the target not chosen by the saccade was turned off. For successful trial, animals had to follow the moving target, remaining within a fixation window of 2 deg. For animal 1, targets moved at 5 deg/s and step size was +1 or -1 deg. Target directions with rightwards (0) and upwards (90). For animal 2, targets moved at 8 deg/s and step size was of -1.6 deg. Target directions with either rightwards (0) and downwards (270) or leftwards (180) and upwards (90). Single and two target stimulus conditions were interleaved. Successful trials were rewarded with marshmallow juice.

Saccade onset and offsets were manually marked using JMWork. Eye velocity was computed from the eye position trace which was collected at 1 kHz and median filtered over 25 ms. The filtered position trace was then differentiated at 25 and 50 Hz and the mean was used as velocity estimate (Mitchell et al., 2015; Pattadkal et al., 2023a).. Trials with saccade latencies within 50-600 ms after motion onset were considered for further analysis. Pre and post-saccadic eye velocities were computed per trial from a 30 ms windows before and after the saccade. The pre-saccadic period ended 25 ms before the actual saccade onset. The post-saccadic period started 25 ms after the actual saccade offset. Post-saccadic eye positions were computer per trial from a 10 ms windows after the saccade, starting 25 ms after the actual saccade offset.

Saccade weights were computed by using the following equation:

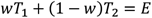

Where w is the saccade weight. T1 and T2 represent the target position vectors for the two target at time of the saccade. E is the eye position vector for post-saccadic eye position.

Pursuit weights were computed by using the following equation:

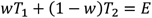

Where w is the pursuit weight. T1 and T2 represent the average pursuit velocity during single target pursuit of target 1 or 2 respectively. E is the eye velocity vector from the trial. For pre-saccadic pursuit weight, T1, T2 and E are from pre-saccadic eye velocities. For post-saccadic pursuit weight, T1, T2 and E are from post-saccadic eye velocities.

To compute the relation between pursuit and saccade weights, a linear fit of the following form was applied:

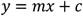

Where y is the pursuit weight, x is the saccade weight, m is the slope of the fit and c is the y-intercept.

Choice probability was estimated by measuring the area under the ROC curve (Green and Swets, 1974). Briefly, pre or post-saccadic pursuit weights were divided into two distributions based on the saccade choice for each trial. The separability of the two distributions was measured by varying the criterion. The hit rate and false alarms obtained for each criterion were used to make the ROC curve. To compute the error bars over choice probability values shown in fig 5, we sampled the weights with replacement 1000 times and used the resulting choice probability distribution to compute the 95% confidence intervals.

## RESULTS

### Post saccadic enhancement of single target pursuit

We first examined the relationship between saccadic and pursuit eye movements when animals tracked a single target using the step-ramp pursuit task (Fig. 1). We are interested in how the presence of a saccadic eye movement at the initiation of pursuit alters the pursuit eye velocity that immediately follows the saccade. If saccadic eye movements reflect an engagement of the pursuit system we expect that post-saccadic pursuit will have a higher gain than pre-saccadic pursuit, and a higher gain than pursuit trials in which no saccade had been made (Lisberger, 1998). Such a comparison requires variability in the latency of the initial saccade during the pursuit trials, which the pursuit behavior provides naturally, or could be enhanced by changing the initial position of the moving target. The task began with the animal fixating at a target at the center of the screen after which a target was stepped either in the direction (positive step) or opposite to the direction of target motion and then the target moved (Fig. 1A). We used both step configurations for animal 1 and only negative steps for animal 2. Both animals successfully tracked the moving target along different directions using a combination of saccades and pursuit (Fig. 1B-D). Pre-saccadic pursuit is generally slower whereas once the animal makes a saccade to the moving target the eye velocity increases to match target velocity (Fig. 1B-D).

**Figure 1:**
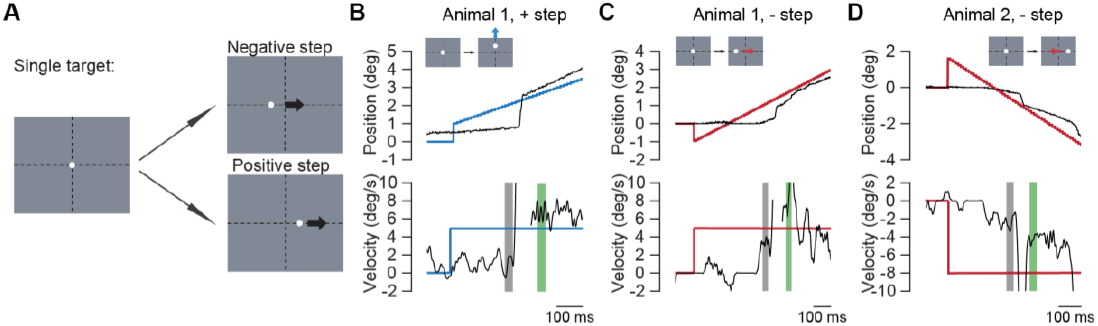
Single-target pursuit task examples A) Shows the scheme for single target step-ramp pursuit task. Animals begin by fixating at a target at center of the screen. The target then makes either a positive or negative step and starts to move. B-C) Example pursuit traces from animals. Top row is showing position trace in the direction of target movement. Bottom row is showing velocity trace in the direction of target movement. Black lines indicate the eye traces. Colored blue and red lines indicate the target motion, along vertical (blue) or horizontal (red) direction. Inset schematics show which direction the target movement is in for each example. Shaded regions indicate the 30 ms windows used to compute pre-saccadic (gray) and post-saccadic (green) pursuit velocity. B is a trace from animal 1, for positive step target moving upwards at 5 deg/s. C is a trace from animal 1, for negative step target moving rightwards at 5 deg/s. D is a trace from animal 2, for negative step target moving leftwards at 8 deg/s.

To quantify the effect of saccades on the initiation of pursuit eye movements we examined the eye velocity signals before and after the first saccade in each pursuit trial. We used an Eyelink 1000 video eye tracker to measure eye position and we observe a systematic perturbation in the eye position traces near the edge of the saccadic eye movements. The eye appears to stop or reverses direction at the edge of the saccade, as others have also noted (Kimmel et al., 2012). These distortions, which may be due to changes in the shape of the pupil during the saccade, could potentially disrupt our measurements of pre- and post-saccadic pursuit. We therefore did not examine pursuit within 25 ms of saccadic eye movements.

To measure the effect of the saccade on pursuit gain we first aligned pursuit trials on the timing of the first saccade and averaged the measured eye velocity that preceded and followed the saccade. On average eye velocity increased in target direction following saccade relative to the eye velocity in same direction preceding the saccade. This effect existed across animals, across step sizes, across target motion directions and across target velocities. (Fig. 2A-C, table 1)

**Table 1:**
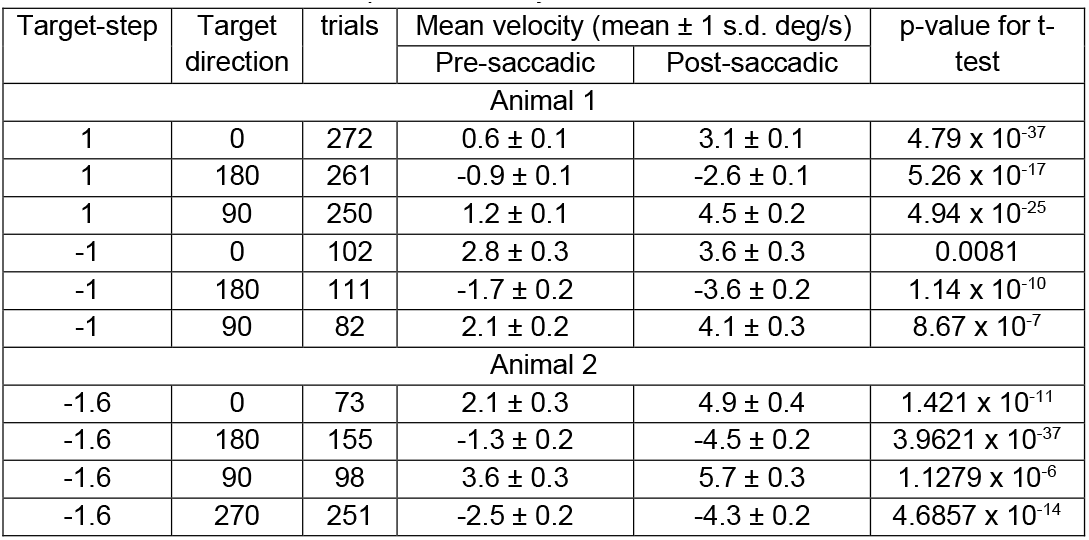
Effect of saccade on pursuit velocity.

**Figure 2:**
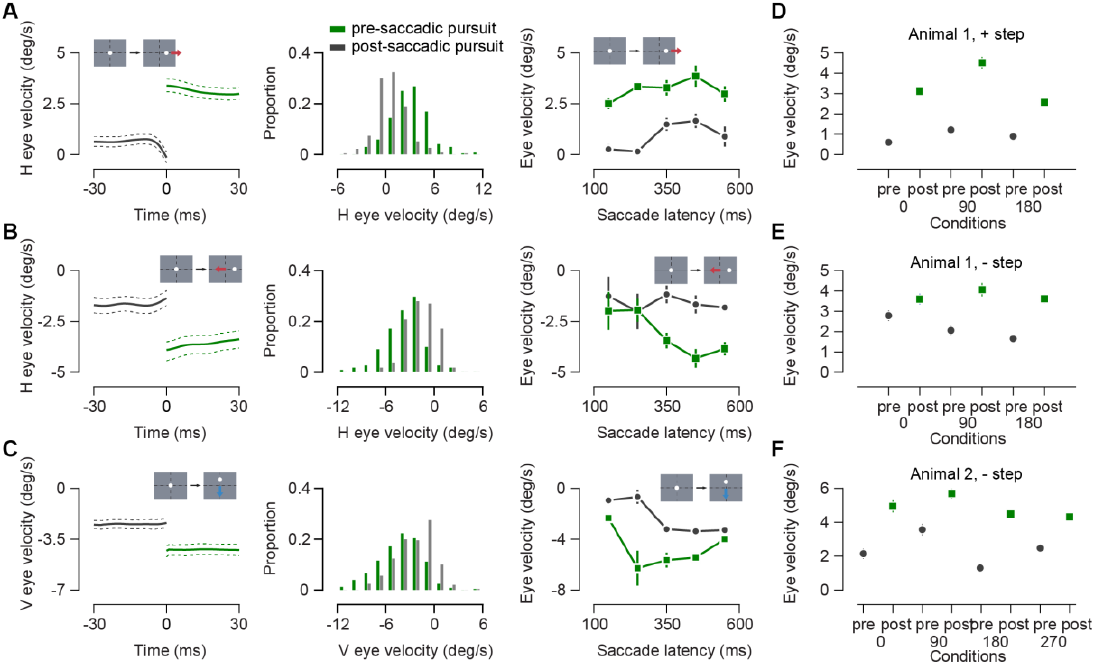
Post-saccadic enhancement of single target pursuit A) Analysis from animal 1 for positive step ramps. First panel shows average eye velocity before (gray) and after (green) the saccade for rightwards target motion. Dotted lines indicate standard error limits. Inset shows the target motion. Second panel shows the distribution of pre and post-saccadic eye velocity for all rightwards single target motion trials. Green indicates post-saccadic pursuit and gray indicates pre-saccadic pursuit. Third panel shows mean velocities in the direction of target motion for different target directions for pre (gray) and post-saccadic (green) pursuit. Error bars are 1 s.e.m. B) Same as A for animal 1, negative step, example motion is leftwards. C) Same as A for animal 2, negative step, example motion is downwards. D) Compares pre (gray) and post (green) eye velocities in trials binned by saccade latencies. Top row is animal 1, positive step, rightwards motion. Middle row is animal 1, negative step, leftwards motion. Bottom row is animal 2, negative step, downwards motion.

This increase in eye velocity could be due to the natural increase in gain that comes at the initiation of pursuit (Schwartz and Lisberger, 1994). That is, while we observe an increase related to the saccade, we already expect an increase to occur because the pursuit system has not yet matched the velocity of the target motion. To distinguish between the effects of time and those of the saccade itself on pursuit velocity, we binned trials based on the timing of the initial saccade and compared the pre- and post-saccadic amplitudes (Fig. 2D-F). Velocities increased for trials with late saccade latencies for both pre and post-saccadic pursuit. But for trials with similar saccade latencies, the post-saccadic pursuit velocity exceeded pre-saccadic pursuit velocity for most bins. Therefore, the rise in pursuit velocity in post-saccadic period is not due to a simple progression in time during the trial, but an effect of the saccade itself. As in other primates, marmoset pursuit gain is increased by the occurrence of a saccadic eye movement.

### Pursuit saccade interaction in two target task

We have demonstrated that for a single target pursuit gain is linked to saccadic eye movements. In the natural world, however, there may be multiple moving objects moving in distinct directions, and animals need to select which target to track. Target selection may engage a combination of saccades and pursuit to foveate the selected target and to stabilize the target motion on the retina. In macaques, a complex interplay between saccades and pursuit has been revealed by employing a task in which two targets are presented and moved, and the animal is given the option to track either target for a reward (Gardner and Lisberger, 2001). As in the single target case, it was found that an increase in pursuit of a target is linked to the saccadic eye movement, but also the direction of the pursuit changed following a saccade. Prior to an initial saccade pursuit eye movements moved in a direction intermediate between the targets whereas after the saccade pursuit moved in the direction of the selected target.

We examined whether there is a similar linkage between saccades and pursuit in marmosets in this two-target task. Animals began with a fixation at the center of screen. Two targets then appeared and moved in different directions at the same speed following a target step. The animal was free to track either target and the other target was extinguished following a variable period (Gardner and Lisberger, 2001) (Fig. 3A). To determine the linkage between the saccadic selection and smooth pursuit we compared the pre-saccadic and post-saccadic pursuit velocity. If saccadic selection is linked to pursuit selection, then we would expect that eye movements following the saccade would track the velocity of the saccadic target, and not of the non-selected target. Indeed, we find that post-saccadic pursuit velocity increases as in the single target case and this increase is towards the target selected by the saccade (Fig. 3B-E).

**Figure 3:**
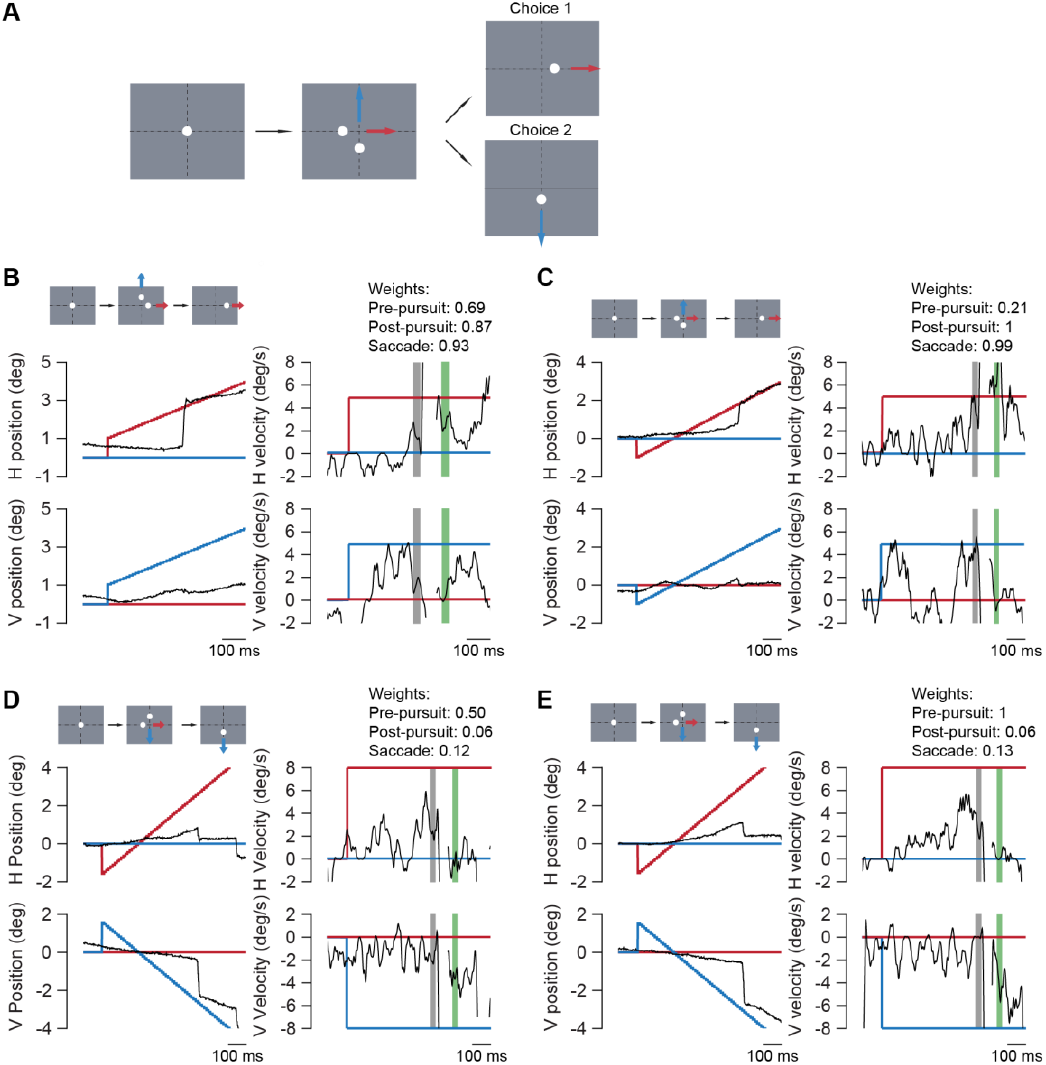
Two-target pursuit task examples A) Shows the scheme for two target step-ramp pursuit task. Animals begin by fixating at a target at center of the screen. The targets then make either a positive or negative step (only negative step is shown) and start to move. After a variable interval, unselected target disappears. B-E) Example pursuit traces from animals. Motion of the target moving horizontally is shown in red and motion of target moving vertically is shown in blue. Eye movement is shown in black. Shaded regions in velocity plots indicate windows used to compute pre (gray) and post-saccadic (green) eye velocities. Target steps and choice is indicated in schematic for each example. Weights for each example are provided. B and C are from animal 1, C and D are from animal 2.

To quantify the evolution of choice during pursuit, we used weights to assign a degree of selectivity in pursuit and of the saccade. Saccade weights are computed by measuring proximity of saccade end point to either of the targets, 0.5 weight being equidistant from both targets and weight of 0 or 1 indicating saccade to either of the targets. Animals used saccades to land closer to one of the targets (see spread of eye position before and after a saccade in Fig. 4A). Pursuit weights were computed in a similar fashion. If the two-target pursuit were the average of two single target pursuits, the pursuit weight would be close to 0.5, whereas a bias towards pursuit of either target shifts the weight towards 0 or 1. For each trial we computed the pre-saccadic pursuit weight, the saccade weight and the post-saccadic weight to determine the interaction between saccades and pursuit.

**Figure 4:**
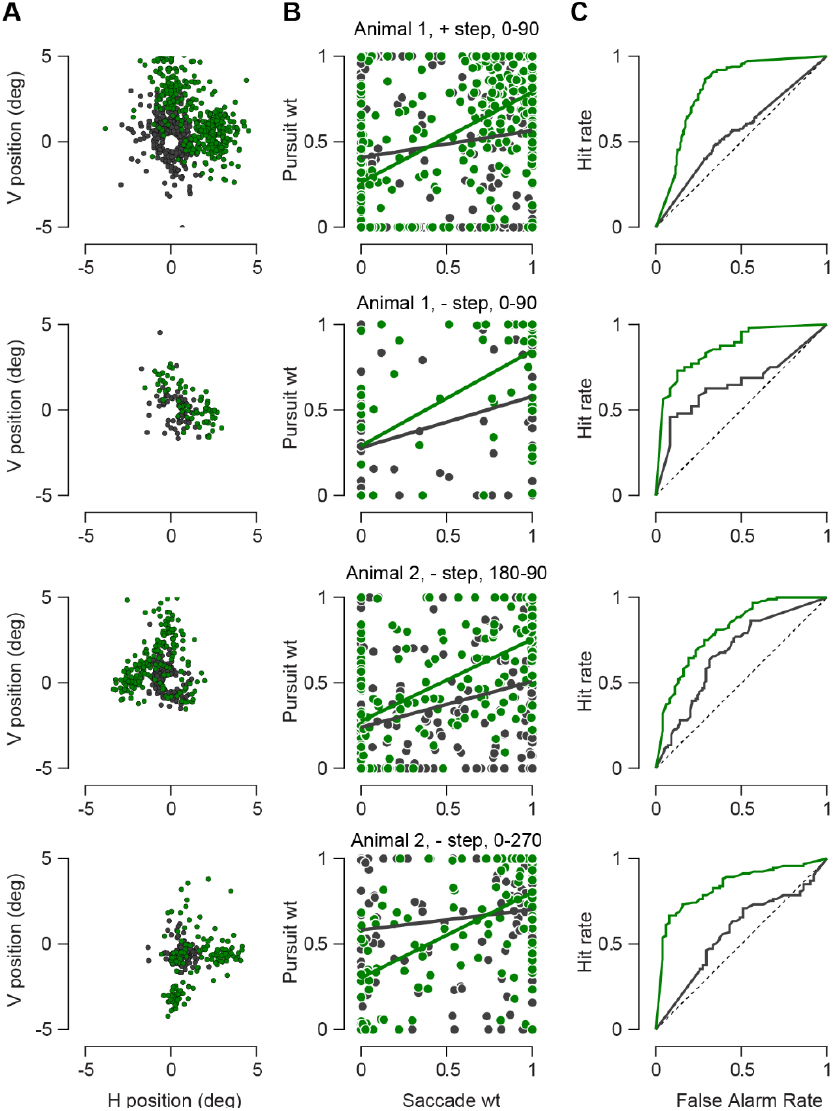
Post-saccadic enhancement of selected target pursuit. A) Shows pre (gray) and post-saccadic (green) eye positions for trials within each condition. B) Compares pre (gray) and post-saccadic (green) pursuit weights to saccade weights. Thick lines indicate the fitted linear regression lines. C) ROC curves for pre (gray) and post-saccadic pursuit weights. Top row in all columns is data for animal 1, positive steps, targets are moving along 0 and 90 degrees. Second row in all columns is data for animal 1, negative step, targets are moving along 0 and 90 degrees. Third row in all columns is data for animal 2, negative step, targets are moving along 180 and 90 degrees. Fourth row in all columns is data for animal 2, negative step, targets are moving along 0 and 270 degrees.

**Figure 5:**
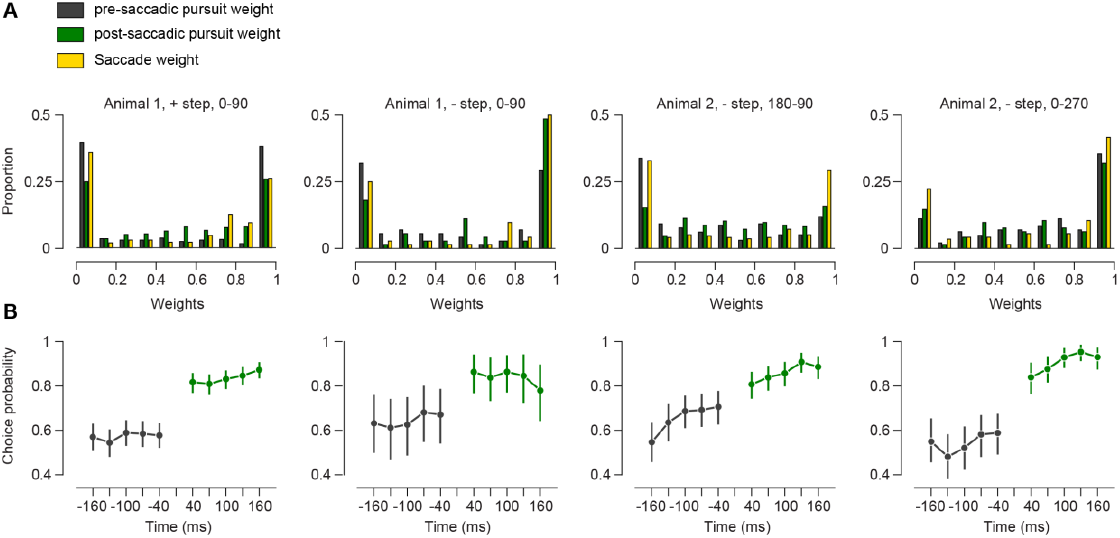
Target selection evolution in time and relation to saccade A) Weight distributions for pre-saccadic pursuit (gray), post-saccadic pursuit (green) and saccades (yellow). Animal, step, and direction conditions are indicated above the histograms. B) Choice probability in different intervals surrounding the saccade. Error bars are 95% confidence intervals on choice probability.

We first tested if saccadic selection influenced pursuit selection. A link between the two would cause post-saccadic pursuit weight to be more related to saccade weight than pre-saccadic pursuit. To quantify this, we measured the slope of the relationship between pre- and post-saccadic pursuit weights and saccade weights. We found that the post-saccadic pursuit weight was more strongly related to the saccade weight than the pre-saccadic pursuit weight (Fig. 4B, table 2). This analysis indicates that post-saccadic pursuit is stronger towards the target that was selected by the saccadic eye movement.

**Table 2:** Summary of relation between pursuit and saccade weights.

| Target-step | Target directions | Trials | Pre-saccadic relation |  |  | Post-saccadic relation |  |  |
| --- | --- | --- | --- | --- | --- | --- | --- | --- |
|  |  |  | Slope | y-intercept | Choice probability | Slope | y-intercept | Choice probability |
| Animal 1 |  |  |  |  |  |  |  |  |
| 1 | 0, 90 | 384 | 0.16 | 0.41 | 0.56 | 0.53 | 0.26 | 0.82 |
| -1 | 0, 90 | 72 | 0.30 | 0.28 | 0.65 | 0.56 | 0.29 | 0.86 |
| Animal 2 |  |  |  |  |  |  |  |  |
| -1.6 | 0, 270 | 144 | 0.12 | 0.58 | 0.67 | 0.50 | 0.30 | 0.81 |
| -1.6 | 180, 90 | 195 | 0.27 | 0.24 | 0.57 | 0.48 | 0.28 | 0.83 |

An alternative way to demonstrate the linkage between saccades and pursuit is to measure the choice probability between the saccadic eye movement and the post-saccadic pursuit. This reflects the degree to which pursuit is predictive of the saccade choice. We first measured how well we could detect the saccade choice using the pursuit weights for pre and post-saccadic pursuit using ROC curves. The post-saccadic pursuit ROC curve is quite convex, reflecting a strong match between the saccadic and pursuit systems following saccade. From this ROC analysis we measured the area under the curve, or choice probability. For both animals, we find that choice probability systematically increases for post-saccadic pursuit (Fig. 4C).

### Is selection absent in pursuit prior to the saccade?

We have demonstrated that pursuit gain increases following saccades and that the direction of pursuit matches the saccade choice post-saccade. But we also find evidence that the pre-saccadic pursuit contains information about the target choice. There are three analyses that indicate that pre-saccadic pursuit exhibits some selection. First, the relationship between the pre-saccadic weight and the saccadic weight is positive (Fig. 4B). This indicates that the pursuit is already biased toward one target. Second, in a similar fashion, the choice probability between pre-saccadic pursuit and the saccade is significantly above 0.5, indicating that there is information in the pursuit eye movements that predicts which target that will be selected. Finally, the distribution of weights reveals that pre-saccadic pursuit also exhibits some selection (Fig. 5A). If the pre-saccadic pursuit were performing a vector average computation, those weights would lie near 0.5. But instead, the pre-saccadic pursuit weights form a bimodal distribution suggesting that the pursuit system has already selected a target. The relatively weak relationship between the pre-saccadic pursuit and saccade weights indicates that it is not uncommon for the saccades to select a different target than the pursuit target. This can be observed in some trials in which saccadic choice did not match the pre-saccadic pursuit choice (ex. fig 3C and E). To study how far back in time pre-saccadic pursuit is indicative of upcoming saccade choice, we computed choice probabilities at different intervals surrounding the saccade (Fig. 5B). The choice probability increased slightly in the interval leading up to the saccade, but the largest change is observed following the saccade.

### Does the saccadic gain enhancement depend on a match to pre-saccade pursuit choice?

Because it is apparent that for some trials the target selected by the pursuit system and the saccadic system are different, we wondered whether changing the selected target would impact the resulting pursuit. If increased gain towards the selected target was affected by a lack of match between saccade and pre-pursuit choice, it would be reflected in the post-saccadic pursuit velocity. To test this idea, we compared post-saccadic pursuit velocities within each saccade choice. For each saccade choice, we divided the post-saccadic velocities into two groups based on whether the saccade choice was matched or mis-matched to the pre-saccadic pursuit choice. We then compared these two groups of post-saccadic velocities (Fig. 6). We do not observe any differences in post-saccadic velocity in the matched vs unmatched groups (KS test, comparisons were made in cases with sufficient trials per subgroup). This indicates that target selective gain enhancement post-pursuit is not dependent on a match with pre-pursuit choice.

**Figure 6:**
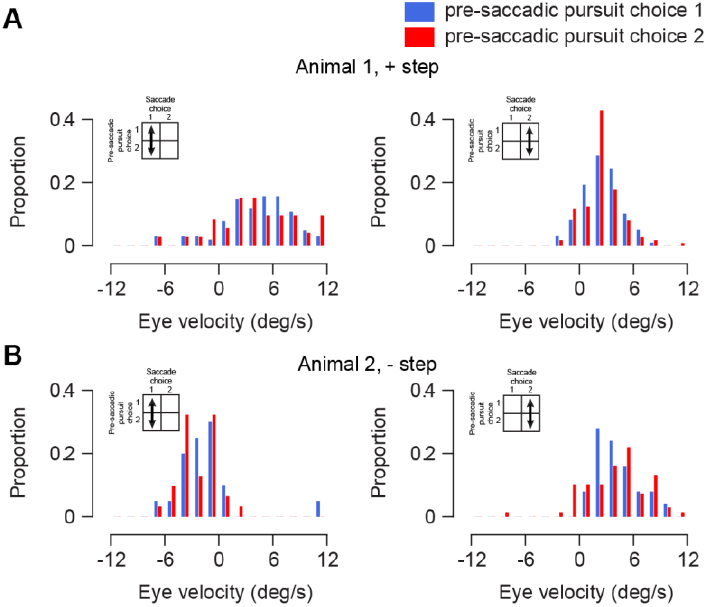
Pre-saccadic pursuit bias does not affect post-saccadic pursuit enhancement A-B) Distribution of post-saccadic eye velocity in the target direction. Left column represents trials where the saccade choice was target 1, right column is for trials with target choice 2. Blue and red bars indicate trials where the pre-saccadic pursuit choice was target 1 (blue) and target 2 (red). A is data from animal 1, positive step condition. B is data from animal 2, negative step, 0-270 condition.

## DISCUSSION

We demonstrate post-saccadic enhancement of smooth pursuit eye movements in marmosets. The post-saccadic eye velocity is higher than pre-saccadic eye velocity irrespective of the initial target step and the saccade latency. When the animal was presented with two target choices to pursue, the choice probability for velocities following a saccade was higher than eye velocities before the saccade. Thus, target selection for pursuit is related in time with target selection for saccade. While the results from the single target task show that there is pursuit enhancement following a saccade, results from the two-target task show that this enhancement is spatially selective and applies only to the selected target.

### Comparison to macaques

As in the macaques (Lisberger, 1998), we find that the eye velocity at a given time was higher if it was post-saccadic compared to pre-saccadic (fig 2). Post-saccadic enhancement occurred independent of the direction of initial target step, whether in same or opposite direction of target motion. The average gains of post-saccadic pursuit were 0.68 (animal 1, positive step, 5 deg/s motion), 0.75 (animal 1, negative step, 5 deg/s motion) and 0.61 (animal 2, negative step, 8 deg/s motion). These gains are similar to previously reported the closed loop pursuit gain in marmosets (Mitchell et al., 2015).

Previous work in macaques shows that when animals are presented with two moving targets, pursuit initiation prior to target choice and in the absence of attentional biases is in the vector average direction (Lisberger and Ferrera, 1997). Once the target is chosen, however, pursuit of that target is not affected by the other target motion. Pursuit choice then follows a winner take all scheme (Ferrera and Lisberger, 1995). When macaques were presented with a task like one in this study, pre-saccadic pursuit represented a vector average between the two target motions and pursuit following saccade was selective for the chosen target (Gardner and Lisberger, 2001). This transition from vector-average to winner-take-all pursuit was linked to the saccade: the pre-saccadic choice probabilities were close to 0.5, indicating vector-average and there is a sharp transition to choice probabilities close to 1 after saccade. In comparison, the pre-saccadic choice probabilities in marmosets were higher than 0.5, indicating that there is already some choice information present in the pre-saccadic pursuit. One reason for this difference may be the differences in the velocities used (20 deg/s in macaque study vs 5 or 8 deg/s in marmosets). The differences in target speeds may affect the saccade latencies (Gellman and Carl, 1991). The mean saccade latency in the macaque study was 224 ms ± 55 whereas mean saccade latency in the two target task in our study is 286 ms ± 105 s.d. (animal 1, positive step), 471 ms ± 112 s.d. (animal 1, negative step), 338 ms ± 158 s.d. (animal 2, negative step). Smaller speeds can also delay pursuit onset though(Ferrera and Lisberger, 1995). The later saccade latency in the present study may give time for the pursuit system to begin to exhibit signatures of target selection.

It seems efficient that the target selection decisions for the saccade system and pursuit system should be shared as they are both have a common goal to view a target (Orban De Xivry and Lefèvre, 2007). While the selection may itself be shared between the pursuit and saccade systems (Gardner and Lisberger, 2001, 2002; Liston and Krauzlis, 2003, 2005), which of the two systems reflects this selection first varies. Some studies (Gardner and Lisberger, 2001, 2002; Schoppik and Lisberger, 2006), argue for a serial linkage between saccadic and pursuit target selection with saccades (either natural or evoked using microstimulation in FEF or SC) capable of triggering selection in pursuit. Other studies (Liston and Krauzlis, 2003, 2005; Case and Ferrera, 2007), show that pursuit preceding saccade can have a different selected target than the saccadic target, though pursuit changes its target to match the saccadic selection around time of saccade. This shift in pursuit can occur prior to the saccade itself with pursuit being able to predict the saccadic target at least 60 ms before the saccade (Liston and Krauzlis, 2003). Coordination dynamics between smooth pursuit and saccade can also depend on the task structure (Erkelens, 2006). The choice of target may be conveyed in parallel to the saccade and pursuit system but the execution of the following saccade and pursuit may occur independently with the time depending on the nature of the task. It is also important to note that a link between saccade and pursuit does not imply that saccades are necessary for pursuit target selection (Ferrera and Lisberger, 1995; Lisberger and Ferrera, 1997; Krauzlis et al., 1999; Garbutt and Lisberger, 2006). However, when both occur in concert, they track the same object. While pursuit selection may not always result in same saccade selection, selection by saccades alters or determines the pursuit target to match. This may be because saccades are the primary means of directing gaze to overtly attend targets.

### Significance of pursuit saccade interaction

Post-saccadic enhancement is known to occur for a variety of eye movements like smooth pursuit in macaques (Lisberger, 1998), ocular following (Kawano and Miles, 1986), disparity vergence (Busettini et al., 1996), tracking responses in marmosets (Coop et al., 2023).

Unlike other eye movements however, post-saccadic enhancement in smooth pursuit is not because of image motion generated by a saccade, but due to the actual saccade. This was shown to be the case by Lisberger 1998 where he observed no enhancement in pursuit upon simulating a saccade by imposing a similar image motion. Pursuit enhancement following a saccade is thought to be a consequence of turning on of the pursuit gain. This pursuit gain is low during visual fixation (Goldreich et al., 1992; Schwartz and Lisberger, 1994) and when switched on, allows for effective tracking of moving target.

Our results demonstrate the similarities in functioning and coordination of eye-movement systems between marmosets and other primates. The accessibility of the underlying neural elements in marmosets (ex:(Mehta et al., 2019; Pattadkal et al., 2023b)) will allow for a circuit level dissection of this cognitive process.

## ACKNOWLEDGEMENTS

Supported by a grant from the National Institutes of Health (R01NS120562).

